# Graded and bidirectional control of real-time reach kinematics by the cerebellum

**DOI:** 10.1101/330555

**Authors:** Matthew I. Becker, Abigail L. Person

## Abstract

The rules governing the relationship between cerebellar output and movement production remain unknown despite the well-recognized importance of the cerebellum in motor learning and precision. In this study, we investigated how cerebellar output sculpts reach behavior in mice by manipulating neural activity in the anterior interposed nucleus (IntA) in closed-loop with ongoing behavior. Optogenetic modulation of cerebellar output revealed monotonically graded and bidirectional control of real-time reach velocity by IntA. Furthermore, kinematic effects were relatively context invariant, suggesting that cerebellar output summates with ongoing motor commands generated elsewhere throughout the reaching movement. These results characterize the relationship between cerebellar output modulation and reach behavior as a bidirectional and scalable kinematic command signal. Our findings illustrate how learned, predictive coding in the cerebellar cortex could be actuated through the cerebellar nuclei to contribute in real time to purposive motor control.

## Introduction

Precise and purposive movement, such as reaching to grasp an object, is controlled by coordinated neural activity across multiple brain regions. When focally disrupted, these neural circuits can produce specific and often debilitating effects on movement. For example, cerebellar damage in humans often leads to reaching movements that over- and under-shoot the target, termed dysmetria, even though the general ability to reach remains intact(Holmes, 1922). While this observation points to the cerebellum as playing a modulatory role over reaching movements, a mechanistic account for these impairments is unresolved.

Research into cerebellar function during movement has largely focused on neural recordings in the cerebellar cortex, where strong correlations exist between neural activity and movement kinematics. For example, populations of Purkinje cells in the cerebellar vermis were recently shown to precisely encode the velocity of saccadic eye movements approximately 20 ms in the future (Herzfeld et al., 2015). For more complex movements such as purposive reach, Purkinje cell populations also reflect movement kinematics (Coltz et al., 1999; Pasalar et al., 2006). Consequently, across a wide range of behaviors, including lower-dimensional behaviors such as eyelid closure (Heiney et al., 2014a) and whisking (Chen et al., 2016), Purkinje neuron encoding of movement kinematics is well established. Purkinje kinematic encoding implies direct and ongoing control of movement, yet the clinical findings of cerebellar dysfunction (e.g. dysmetria) resemble deficits in precision and/or coordination, as opposed to movement generation. These observations raise a fundamental question: what does the cerebellum contribute to ongoing movement?

Determining the function of cerebellar kinematic encoding requires understanding the rules governing the translation of cerebellar output into motor control. Purkinje neurons converge in the cerebellar nuclei, which as the sole output structures of the cerebellum, are a critical link to the rest of the motor system. In tests of cerebellar function during reaching movements, pharmacological inactivations (Milak et al., 1997; Mason et al., 1998; Cooper et al., 2000; Martin et al., 2000) and lesions (Low et al., 2018) of the anterior Interposed Nucleus (IntA) were shown to recapitulate the dysmetria observed in cerebellar patients. However, since these neural activity manipulations last orders-of-magnitude longer than a single reaching movement, differentiating real-time control from related motor functions, such as movement planning or initiation (Meyer-Lohmann et al., 1977), has remained problematic. In other work, electrical or optogenetic stimulation of reach-related cerebellar areas was shown to result in forelimb movement (Rispal-Padel et al., 1982; Witter et al., 2013; Lee et al., 2015), demonstrating the sufficiency of cerebellar output to initiate non-purposive limb movements. The applicability of these results to natural cortically-driven reach behavior (Denny-Brown, 1950; Guo et al., 2015), which is a dynamic movement that evolves through time to a target, remains unclear.

Taken together, previous functional studies of cerebellar contributions to purposive behavior were limited by temporal imprecision and lack of control over the acute behavioral context of neural manipulation. Recent advances in computer vision and optogenetics provide novel leverage to address these limitations, however, and allow tests of IntA function specifically during ongoing reach behavior. IntA projection neurons heavily target motor cortex (via thalamus) and the red nucleus (Eccles et al., 1967; Houck and Person, 2015), both important motor control centers for reach behavior (Houk, 1991; Alstermark and Isa, 2012). This anatomical organization indicates that fluctuations in IntA activity during reach (Burton and Onoda, 1977; Fortier et al., 1989; Goodkin and Thach, 2003) can directly influence ongoing movements generated elsewhere, yet the rules governing this interaction remain uncharacterized.

To test the causal role of cerebellar output in online control of purposive reaching movement, we designed a kinematic closed-loop system that enables brief, scalable optogenetic manipulation tied directly to ongoing reach kinematics in mice. This technology allowed us to characterize the functional relationship between cerebellar output from IntA and movement kinematics, including its sign, strength, and context-dependence. We show that modulation of cerebellar output results in monotonically scalable, bidirectional control of real-time reach velocity. Furthermore, the invariant directionality of the kinematic effect of IntA activity modulation across multiple phases of reach implies that cerebellar output is additively integrated with ongoing motor commands. These results describe a continuous and scalable kinematic command signal in IntA capable of modulating reach behavior in real time. Moreover, our findings suggest that learned predictive coding in the cerebellar cortex has direct and ongoing access to a kinematic motor controller in the cerebellar nuclei.

## Results

### Kinematic closed-loop excitation of cerebellar output affects real-time reach kinematics

Previous electrophysiological experiments implicated the anterior interposed nucleus (IntA) in reaching movements, but causal tests of its role relied on pharmacological circuit manipulations that lasted orders-of-magnitude longer than a single reaching movement (Milak et al., 1997; Mason et al., 1998; Cooper et al., 2000; Martin et al., 2000; Low et al., 2018). Thus, these studies were limited in defining the temporal contribution of cerebellar output in ongoing motor control. Moreover, because reaches evolve over time, it was unclear whether cerebellar output preferentially influenced specific phases of movement. To circumvent these issues, we designed a kinematic closed-loop system that tracks mouse reach kinematics in real time and ties neural manipulation directly to ongoing behavior (Fig.1a). This novel paradigm enabled us to characterize the real-time contribution of cerebellar output to movement by manipulating cerebellar output during a purposive movement on a timescale that is brief relative to the duration of a single reach. We first adapted a motion capture system designed for human kinematic analysis to the scale of mouse reach behavior in order to track the 3D position of a mouse’s paw. Our kinematic tracking system had a spatial resolution of 150 µm and tracking latency of less than 1 ms, resulting in real-time tracking of reach trajectories with high spatiotemporal fidelity (Fig. 1b; Supplementary Video 1). Reach kinematic data was analyzed in real time to trigger optogenetic stimulation at specific, user-defined kinematic landmarks with a 0.5 ms average processing latency (Supplementary Fig. 1). We trained mice on a purposive reach task in which freely-behaving animals retrieve small food pellets from a pedestal (Whishaw, 1996; Azim et al., 2014) (n = 7; Fig. 1 a,b). Mice attained peak success rates of 65 ± 7% over the course of several days of training. Trained reaches lasted 316.3 ± 102.5 ms (outreach; Supplementary Fig. 1), much longer than the total closed-loop latency of 9.5 ms (8 ms interframe interval, 1 ms tracking, 0.5 ms processing). Thus, during a single reaching movement, we could implement temporally precise, short-latency optogenetic stimulation tied directly to ongoing reach kinematics.

**Figure 1.**
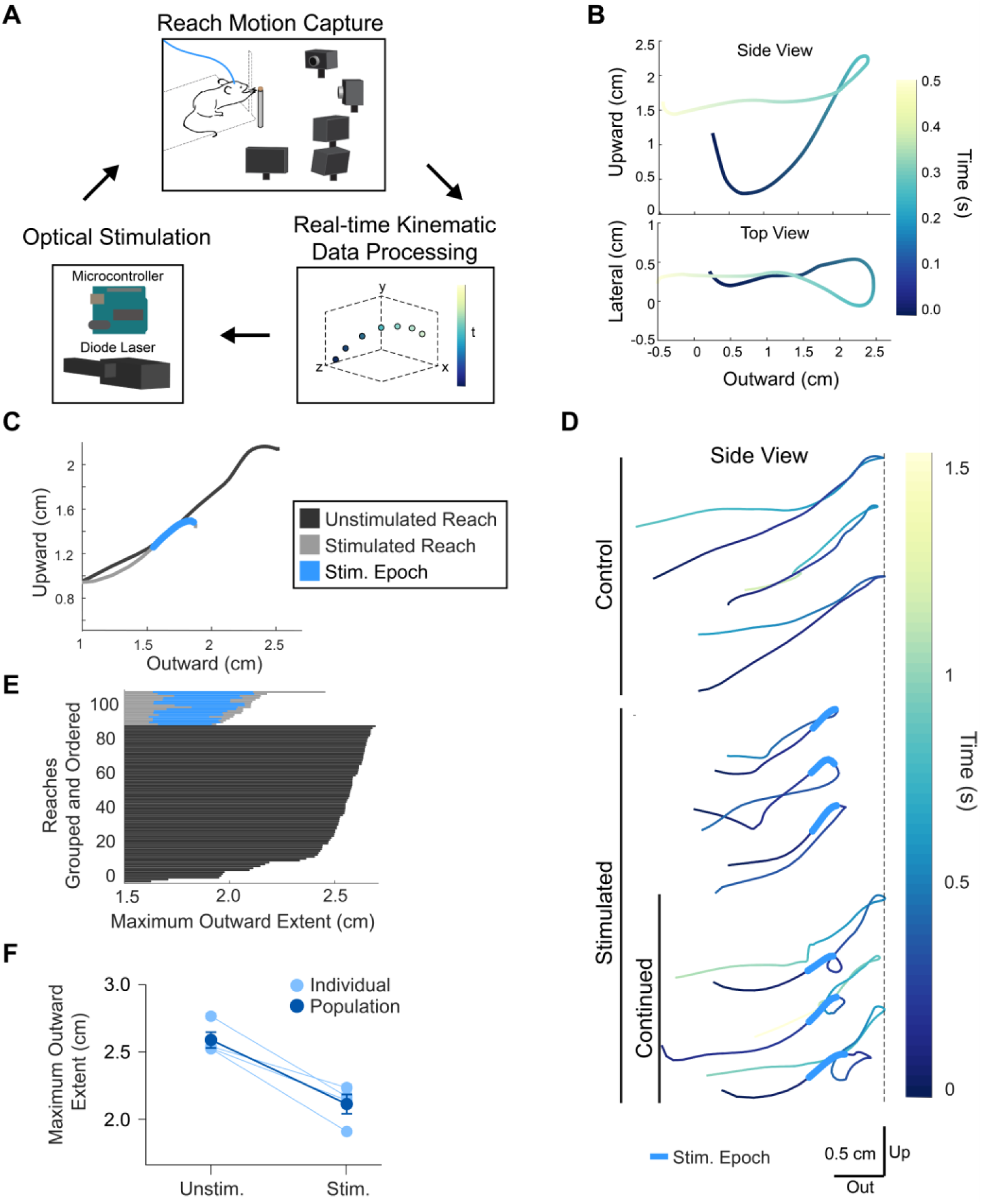
Kinematic closed-loop stimulation of IntA during reach. **A**. Schematic of kinematic closed-loop system, in which real-time tracking of mouse reach kinematics is used to trigger optogenetic stimulation. **B**. Example mouse 3D reach trajectory tracked in real time. ‘Upward’, ‘Lateral’, and ‘Outward’ represent the three spatial dimensions, with ‘Outward’ being in the direction of the food pellet target. **C**. Examples of outward phase of unstimulated and stimulated reach trajectories using kinematic closed-loop optogenetic excitation of IntA (‘ChR2’) at a specific kinematic landmark (see Methods). **D**. Examples of complete unstimulated reach trajectories (top 3) which extend fully to the target location (dashed line); stimulated reach trajectories (middle 3), that return fully after stimulation; and stimulated reaches that continue out toward the target following the initial direction reversal (‘Continued’, bottom 3). **E**. Reach trajectories from a single behavioral session at their initial maximum extent in the Outward direction. Reaches are grouped by type (Unstimulated or Stimulated) and ordered by maximum extent for visual clarity. **F**. Average maximum outward extent of unstimulated and stimulated reaches (4/4 animals, p < 0.0001; mean ± SE).

To test the hypothesis that the cerebellum actively sculpts movement kinematics, we used optogenetic stimulation to manipulate cerebellar output at a specific kinematic landmark relative to typical maximal reach velocity (Supplementary Fig. 1). This positional landmark, a vertical plane located at the opening of the behavior arena that the animals reach through (see Methods), was consistently applied across all animals tested. We injected trained animals with AAV-hSyn-hChR2-mCh (‘ChR2’) and implanted an optical fiber in IntA ipsilateral to the arm used for reaching (Supplementary Fig. 2). When mice reached for the target, we delivered a 50 ms train of blue light pulses (2 ms, 100 Hz) in closed-loop, stimulating on a random 25% of reaches to avoid anticipation.

**Figure 2.**
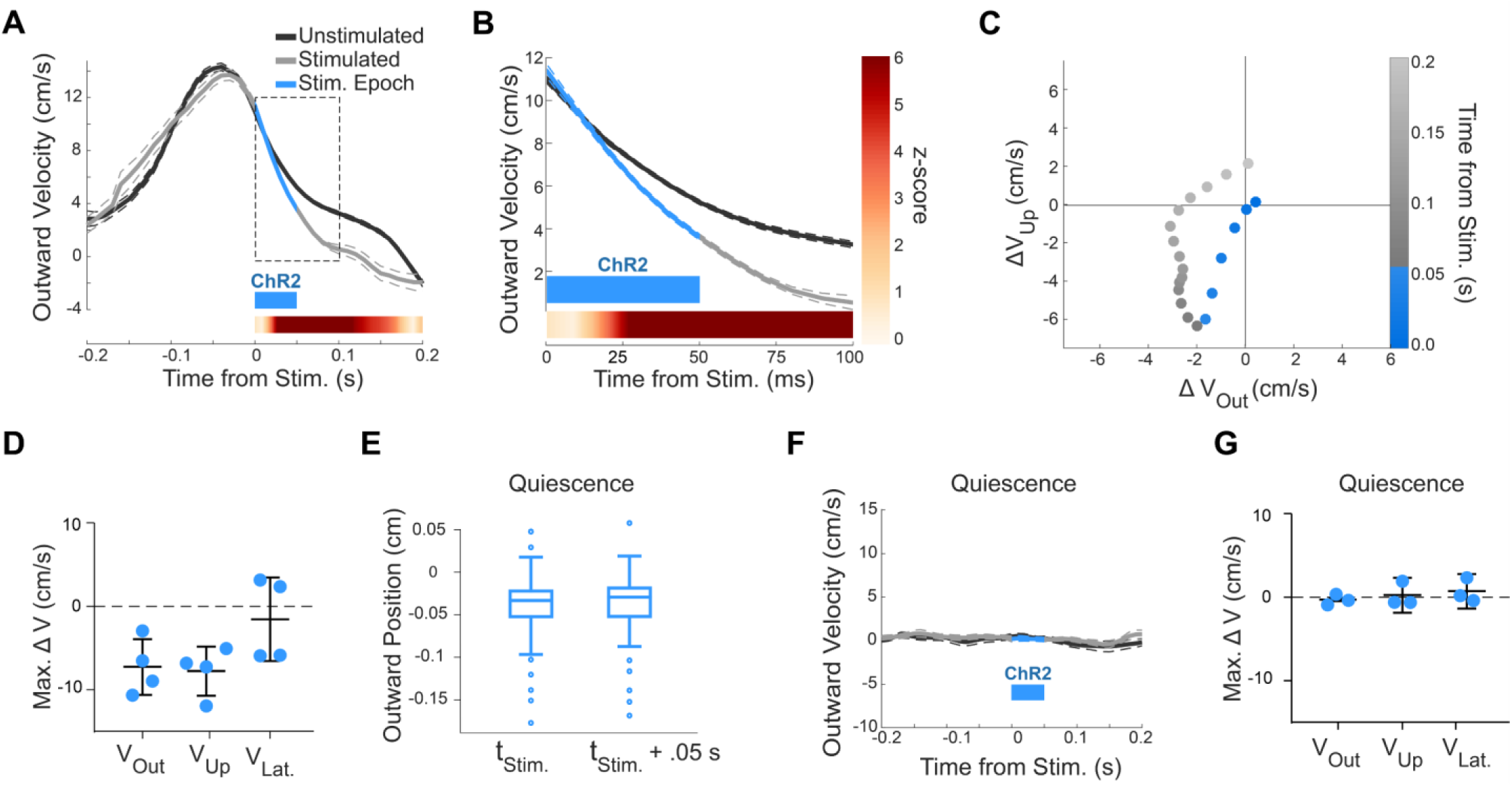
Velocity effects of kinematic closed-loop stimulation of IntA. **A,B.** Example of average outward velocity of unstimulated and stimulated reaches from one animal, aligned to time of stimulation (stimulated reaches) or kinematic landmark (unstimulated reaches). The linearly interpolated z-score (colorbar) shows the timecourse of statistical divergence between the unstimulated and stimulated reach velocities (see Methods). All velocity profiles display mean (solid line) ± SE (dashed lines). **C**. Two-dimensional timecourse of average ChR2-driven reach velocity changes (Stimulated – Unstimulated) from same example shown in **A**. Values at the origin indicate no difference between stimulated and unstimulated reaches. **D**. Summary of average maximum velocity effect magnitudes (Reaching; Stimulated – Unstimulated) in each of the three spatial dimensions across subjects (Outward and Upward directions, 4/4 animals, p < .0001; lines, mean ± SE). **E**. Box-and-whisker plot of the effect of optogenetic stimulation during quiescence and feeding on average paw position (p > 0.05). **F**. Example of the effect of optogenetic stimulation during quiescence and feeding on average paw velocity (p > 0.05). **G**. Summary of average maximum velocity effect magnitudes (Quiescence; Stimulated – Unstimulated; Outward, 3/3 animals, p > 0.05; Upward, 2/3 animals, p > 0.05) in each of the three spatial dimensions across subjects.

Brief excitation of IntA during outreach resulted in clear and consistent kinematic effects. Stimulated reaches showed a short-latency reversal in outward trajectory (latency from stim; mean = 78 ± 22 ms; n = 4), resulting in reaches that did not progress as far towards the target as unstimulated reaches (Fig. 1c; Supplementary Videos 2-4). While some reaches were truncated in the outward direction after stimulation, many reaches continued toward the target after the initial direction reversal, demonstrating active behavioral engagement throughout and beyond the stimulation epoch (latency from stim; mean = 211 ± 33 ms; n = 4 mice; ‘Continued reaches’ Fig. 1d; Supplementary Video 3). Across animals, an average of 75 ± 9% of stimulated reaches continued out towards the target subsequent to the initial direction reversal. To quantify the positional effect of stimulation, we clipped both unstimulated and stimulated reach data trajectories at their first endpoint in the outward direction (Fig. 1e). Across animals and across days, stimulation consistently resulted in reaches not progressing as far towards the target (4/4 animals, Wilcoxon rank sum, p < 0.0001; Fig. 1f; Supplementary Video 4).

To better understand the temporal and directional properties of the cerebellum’s influence on purposive behavior, we analyzed the effects of IntA optogenetic stimulation on reach velocity. Unstimulated reaches possessed a characteristic ‘bell-shaped’ velocity profile during outreach(Flash and Hogan, 1985) (Fig. 2a, black traces). As implied by the positional effects, excitation resulted in a clear slowing of the paw in the outward direction (Fig. 2a). Average unstimulated and stimulated velocity trajectories in the outward direction were indistinguishable up to the point of stimulation, at which point they quickly diverged as stimulated reaches slowed prematurely (Fig. 2b). The stimulation-induced decrease in outward velocity was rapid, beginning even before termination of the pulse train (Wilcoxon rank sum, z-score > 2.5, mean latency 18 ± 5 ms). Subtraction of the average unstimulated and stimulated velocity trajectories showed a consistent slowing in the outward direction as well as the upward direction (Stim. – Unstim.; Fig. 2c). In all animals tested, excitation of IntA during outreach caused decreases in outward and upward velocity relative to unstimulated reaches (4/4 animals, Wilcoxon rank sum, p < 0.0001; Fig. 2d). This remarkable similarity in the directionality of the kinematic effects across animals implies consistency in the relationship between cerebellar output modulation, specifically from the IntA, and reach kinematics.

These results indicate that brief optogenetic excitation of cerebellar output at a consistent kinematic landmark during outreach has reproducible effects on ongoing reach kinematics. However, the nature of the effect (the paw moving back towards the body) prompted concern that the underlying mechanism was off-target to reach control or related to an aversive response. While the existence of ‘continued’ reaches argues against this concern (Fig. 1d), we also performed control experiments in which the same optogenetic stimulation protocol was applied during periods of non-reach behavior (i.e. feeding and quiescence; see Methods). Stimulation outside of the context of active reaching had little or no effect on paw position or velocity (Fig. 2e-g; Supplementary Video 5; outward direction velocity, 3/3 animals, Wilcoxon rank sum, p > 0.05; upward direction, 2/3 animals, Wilcoxon rank sum, p > 0.05). While we found that stimulation of IntA at higher optical powers is able to cause non-purposive limb movements at rest, consistent with previous observations (Witter et al., 2013; Lee et al., 2015) (data not shown), our results nevertheless demonstrate an increased kinematic sensitivity to cerebellar modulation during active reach. These data position the cerebellum as a kinematic modulator of purposive limb behavior.

### Graded and bidirectional control of reach velocity

If the kinematic control demonstrated above is a mechanism through which the cerebellar nuclei control kinematics with precision, then we would predict that that graded fluctuations of IntA activity would result in proportional changes in reach velocity. We therefore stimulated IntA in mice expressing ChR2 with a range of optical powers, and predicted that lower powers would result in smaller amplitude effects on reach kinematics. As predicted, over the range of light intensities tested, lower optical power caused concomitantly smaller effects on outward and upward velocity (Fig. 3a). As optical power was decreased, the maximum difference between average unstimulated and stimulated velocities decreased in parallel (Stim. – Unstim.; 1.0 mW: -9.4 ± 0.7 cm/s; 0.5 mW: -7.9 ± 0.1 cm/s; 0.25 mW: -7.5 ± 0.2 cm/s; 0.1 mW: -1.3 ± 0.1 cm/s). To rule out the possibility that decreased average effect sizes reflected changes in the frequency of an all-or-none effect rather than truly graded effect magnitudes, we calculated the likelihood of reach velocity values across the reaching epoch for both unstimulated and stimulated reaches (see Methods). As optical power was reduced, a clear merging of the unstimulated and stimulated distributions occurred without any visible presence of an all-or-none effect, demonstrating graded effect sizes across the population of stimulated reaches (Fig. 3b). Next, we examined the pattern of changes in outward and upward reach velocity by subtracting the average velocity profiles of stimulated reaches from unstimulated reaches (Fig. 3c). Interestingly, as the amplitude of velocity effect size decreased, we observed maintenance of the spatial covariance of outward and upward velocity effects, suggesting scalable but clearly directional control of reach velocity by IntA output modulation.

**Figure 3.**
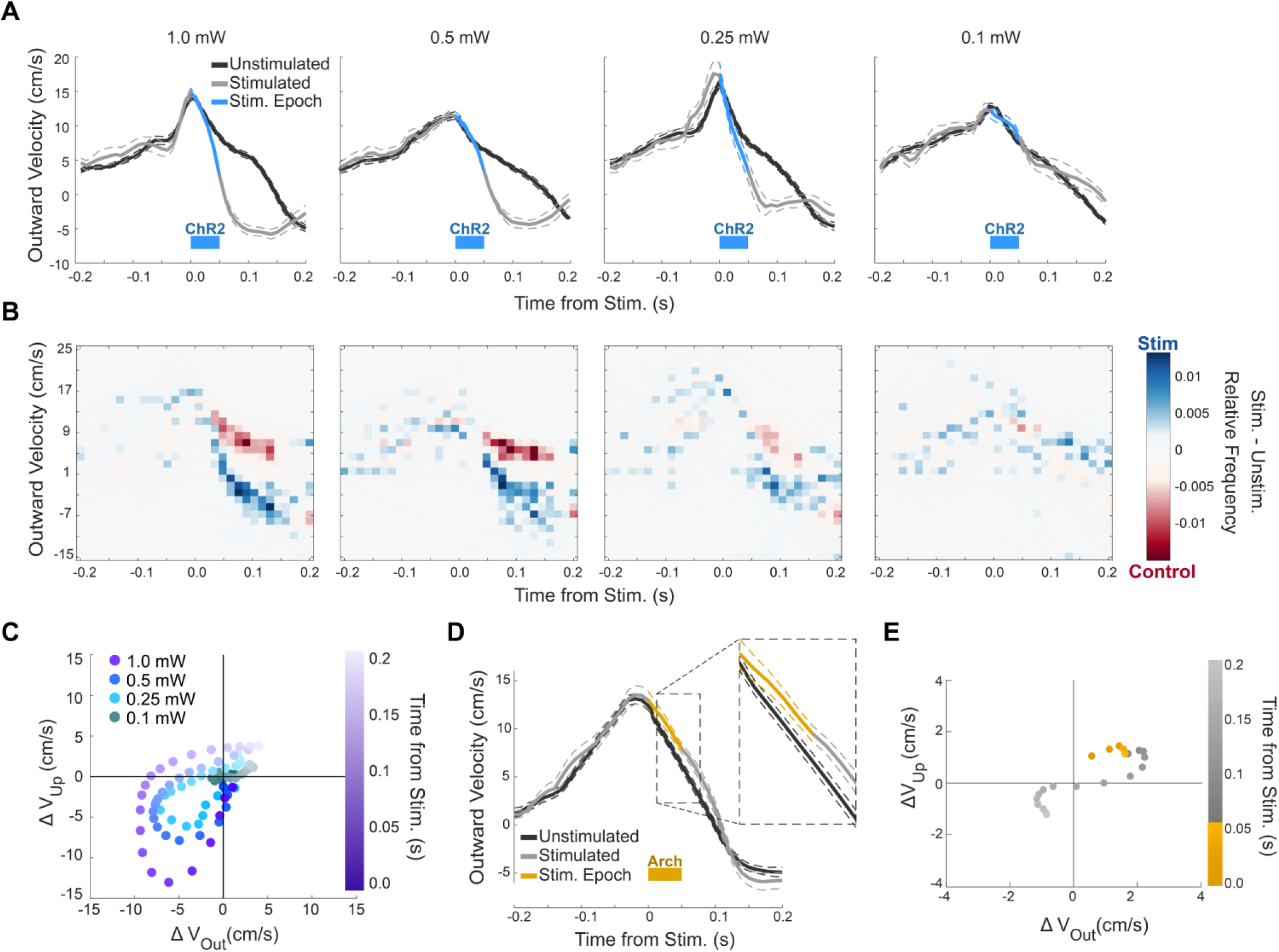
Graded and bidirectional control of reach kinematics by IntA modulation. **A.** Average outward velocity of unstimulated and stimulated reaches with decreasing optical power of optogenetic stimulation (‘ChR2’) from high (1.0 mW, far-left) to low (0.1 mW, far-right). **B.** Heatmap of relative frequency of unstimulated and stimulated reach velocity values. Individual frequency heatmaps for unstimulated reaches and stimulated reaches were normalized and subtracted (Stimulated – Unstimulated) to view the relative prevalence of reach velocity values in each distribution. **C.** Two-dimensional timecourse of average ChR2-driven reach velocity changes (Stimulated – Unstimulated) for each optical power level tested in **A**. **D.** Example of average outward velocity in response to kinematic closed-loop optogenetic inhibition of IntA (‘Arch’). **E.** Two-dimensional timecourse of average Arch-driven reach velocity changes (Arch; Stimulated – Unstimulated) from same example shown in **D**.

Graded levels of IntA excitation resulted in proportional decreases in outward and upward reach velocity, which raised the question of whether the directional relationship between cerebellar output modulation and movement kinematics extended to firing rate decreases. IntA premotor output neurons exhibit tonic baseline firing rates in vivo of ∼20-80 Hz (Armstrong and Edgley, 1984; Rowland and Jaeger, 2005) and receive massive inhibitory input from Purkinje neurons, potentially allowing for a bidirectional dynamic range of kinematic control. We therefore hypothesized that decreased activity of IntA during outreach would result in increased outward and upward velocity, opposite to that seen during excitation. To test this hypothesis, we performed closed-loop inhibition of IntA using the inhibitory opsin Arch (AAV-hSyn-Arch3.0-YFP), which in control recordings strongly inhibited IntA neurons (Supplementary Fig. 3) (Chow et al., 2010; Mattis et al., 2011). Consistent with the idea that IntA exerts bidirectional control of kinematics, we observed opposite kinematic effects in response to inhibition compared to excitation of IntA neurons. Outward reach velocity increased temporarily in response to stimulation (Stim. – Unstim.; 3.3 ± 1.2 cm/s; Fig. 3d). 2-dimensional analysis revealed that both outward and upward velocities were increased in response to inhibition, directionally opposite to the results observed during excitation (Fig. 3e).

The graded kinematic effects of IntA excitation, combined with the opposite directional effects of IntA inhibition, identify a monotonic relationship between IntA activity and velocity control. Over the population of animals tested, we always observed a monotonic relationship between optical power and kinematic effect size, including both increases and decreases of IntA activity (Fig. 4a). Inhibition with Arch consistently resulted in increased outward and upward velocity, while graded levels of excitation with ChR2 always resulted in monotonic effects sizes of decreased outward and upward velocity (Outward direction: 4/4 ChR2 animals, 3/3 Arch animals, Wilcoxon rank sum, p < 0.05; Upward direction: 4/4 ChR2 animals, 2/3 Arch animals, Wilcoxon rank sum, p < 0.05; Fig. 4a,b). Peak kinematic effect sizes of Arch stimulation were smaller in amplitude and longer-latency than those of ChR2 stimulation (ChR2 latency 43 ± 10 ms; Arch latency 67 ± 7 ms; ChR2 magnitude (speed) 10.8 ± 3.2 cm/s; Arch magnitude (speed) 3.0 ± 1.4 cm/s; Fig. 4c). This observation could be due to differences in the dynamic range of the effect of brief (50 ms) excitation and inhibition on IntA firing rates (Supplementary Fig. 3). More specifically, for an IntA neuron firing at 20-80 Hz, complete inhibition for 50 ms would remove on average only 1-2 spikes. Alternatively, ChR2 stimulation could add up to 20 spikes during the 50 ms pulse train, since IntA neurons can fire at rates as high as 400 spikes/s in response to stimulation (Bagnall et al., 2009). In summary, we found a bidirectional, monotonic relationship between IntA activity and real-time control of limb velocity: excitation of IntA decreased outward and upward velocity, while inhibition of IntA increased outward and upward velocity.

**Figure 4.**
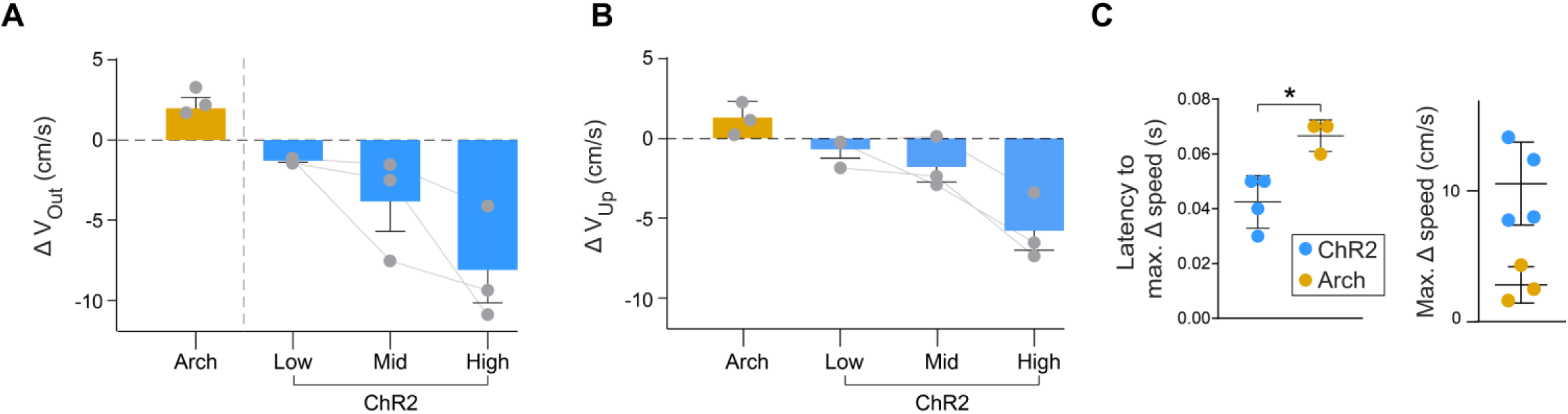
Summary of graded excitation and inhibition of IntA on reach kinematics. **A-B.** Summary of average effect sizes (Stimulated – Unstimulated; mean ± SE) from graded levels of excitation (‘ChR2’) or inhibition (‘Arch’) on reach velocity in the outward (**a**) and upward (**b**) directions across animals. **C.** Latency (left panel) and magnitude (right panel) of maximum effect of stimulation on reach speed under excitation (ChR2) and inhibition (Arch) (lines, mean ± SE).

### State-dependence of cerebellar influence of reach motor commands

The functional relationship identified thus far relates to kinematic effects of cerebellar output modulation at a specific position in reach. However, several lines of evidence point to the possibility that the relationship between IntA activity and movement kinematics might be dependent on motor state (i.e. position and velocity of the limb). For example, we observed that stimulation during non-reach behaviors such as feeding and quiescence did not consistently affect movement kinematics, implying that the integration of cerebellar output with ongoing activity in other brain regions could be highly nonlinear in its net effect on movement. Additionally, the activation of muscle synergies (e.g. via microstimulation of the spinal cord) is known to cause movements that are highly dependent on initial limb state(Mussa-Ivaldi et al., 1994). These results led us to test whether the kinematic effect of IntA modulation is strongly state-dependent within the reach itself, where the general behavioral context (reaching) is constant while the specific state of the limb (position/velocity) is rapidly changing.

Our experimental paradigm afforded the unique opportunity to examine the state-dependence of the relationship between IntA activity modulation and reach kinematics. We added two additional kinematic landmarks flanking our original stimulation location, each comprising a unique initial position and velocity (3/3 animals, Kruskal-Wallis, p < 0.0001; Fig. 5a). We then excited IntA in closed-loop separately at each location (called ‘early,’ ‘middle,’ and ‘late’). Interestingly, we observed similar initial decreases in outward and upward velocity across tested landmarks, regardless of the position or velocity of the paw at the time of stimulation (Fig. 5b,c). However, the magnitude of the difference between average unstimulated and stimulated velocities was not perfectly consistent across the kinematic landmarks tested, as we observed lower average effect magnitudes later in the reach (Early: -7.5 ± 3.0 cm/s; Middle -6.4 ± 3.0 cm/s; Late: -2.8 ± 2.3 cm/s; Friedman Test, p < 0.05; Fig. 5e).

**Figure 5.**
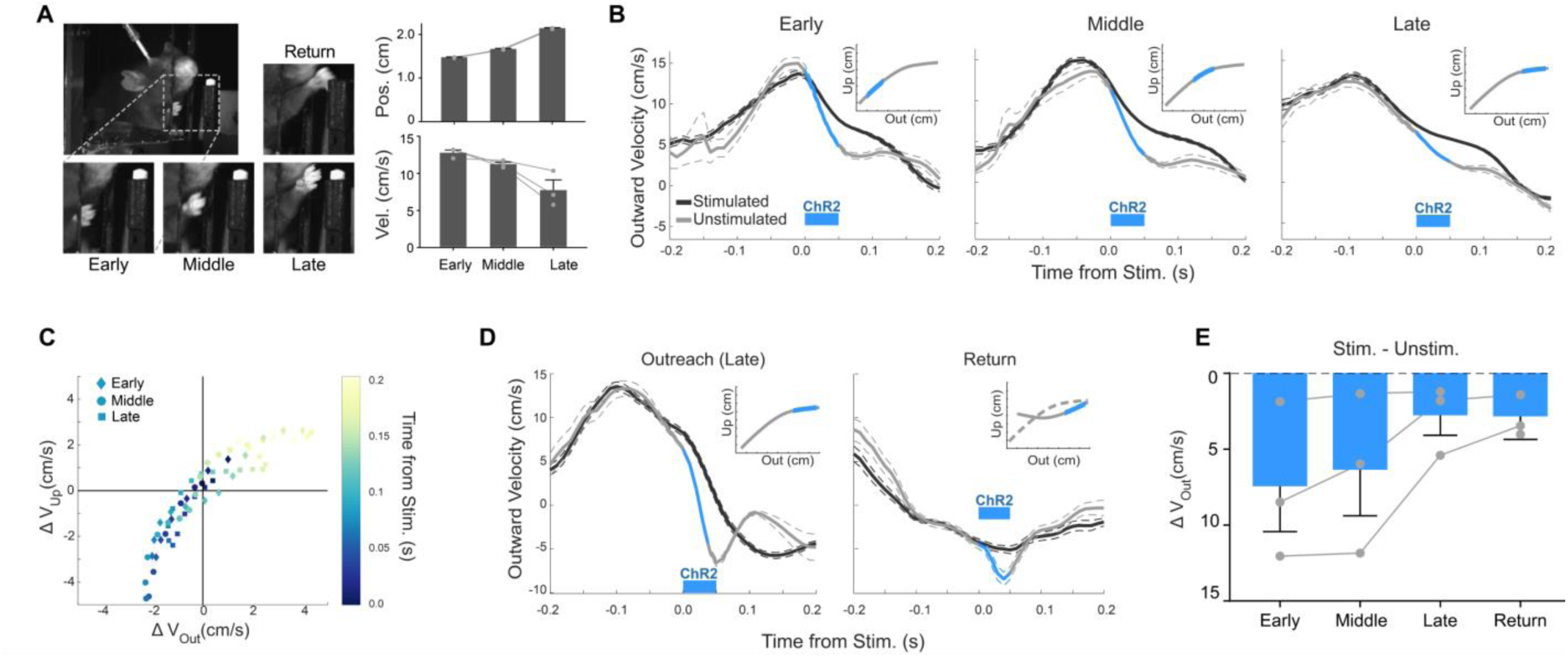
Context dependence of IntA stimulation on reach kinematics. **A.** Images of paw at three kinematic landmarks used to trigger IntA stimulation (left), and the associated positional and velocity characteristics at each landmark (mean ± SE, right). **B.** Average outward velocity of reaches stimulated with ChR2 at different kinematic landmarks overlayed with unstimulated reaches. Insets: schematic of stimulation trigger points. **C.** Two-dimensional timecourse of average ChR2-driven reach velocity changes for three stimulation landmarks. **D.** Average outward velocity of reaches stimulated with ChR2 at ‘Late’ and ‘Return’ landmarks overlayed with unstimulated reaches. The positional landmark triggering ‘Late’ and ‘Return’ stimulation was identical. **E.** Summary of average effect sizes (mean ± SE) on outward velocity from stimulation at each kinematic landmark (‘Early’, ‘Middle’, ‘Late’, and ‘Return’).

One interpretation of this result is that IntA controls reach progression instead of direct control of kinematics. This possibility is consistent with our observation that the kinematic effect of excitation opposes the ongoing direction of motion; for reaching movements that are directed outward and upward, excitation always resulted in decreased outward and upward velocity (see Fig. 2f). In addition, decreased effect sizes later in the reach could be caused by the reach being closer to completion at the later landmarks. Therefore, we designed an experiment to distinguish between control of reach progression and direct control of reach kinematics. We reasoned that if cerebellar output controls reach progression, excitation of IntA during the return phase of reach, after the animal has already obtained the pellet, should cause *increased* outward velocity (in opposition to the current direction of motion). We therefore stimulated on the return phase of the reach at an identical positional landmark to the ‘late’ kinematic landmark used during outreach. In this way, we held position constant but allowed the direction of reach progression to vary. In response to ChR2 stimulation during return, we consistently observed a further decrease in outward velocity (i.e. increase in inward velocity), in which the paw moved back towards the animal faster than in the unstimulated condition (Stim – Unstim; 3/3 animals; Wilcoxon rank sum, p < 0.01; -2.9 ± 1.5 cm/s; Fig. 5d,e). Thus, these results indicate that IntA exerts an invariant directional effect on reach kinematics, in which excitation of IntA moves the paw downward and inward. The consistency of the directional effect, regardless of ongoing movement direction, implies that IntA velocity commands are added onto the ongoing motor command structure. This interpretation is consistent with experimental findings in the eye movement literature, reporting summation of the output of a positional integrator and vestibular signals (Skavenski and Robinson, 1973; Ramachandran and Lisberger, 2006).

## Discussion

In this study, we investigated the functional role of cerebellar output during reach behavior by manipulating neural activity in IntA at specific behavioral contexts and measuring the effects on movement kinematics. Using a novel kinematic closed-loop system, we found that graded, bidirectional manipulations of IntA result in graded, bidirectional effects on reach velocity. The directionality of the kinematic effect was consistent across all animals tested; excitation of IntA decreased outward and upward velocity, while inhibition of IntA increased outward and upward velocity. Moreover, the pattern of kinematic effect was maintained during decreasing magnitudes of stimulation, as the amplitude of the effect concomitantly decreased. In testing the role of IntA at several unique kinematic landmarks throughout reaching movement, the direction of kinematic effect was consistent, implying that cerebellar activity is added onto the underlying motor command structure. These experiments characterize basic properties of the functional relationship between cerebellar output and ongoing movement kinematics during purposive reach behavior. Taken together, our results implicate IntA as a stable kinematic controller capable of actuating learned cerebellar predictions into precise movement.

### Properties of kinematic control by IntA during reach behavior

Two remarkable features of kinematic control by IntA are its consistent directionality and proportional effects on velocity, which position IntA as a stable motor controller for tuning the velocity of purposive limb movements. The surprisingly consistent directionality of kinematic effect across animals suggests that hard-wired anatomical projections downstream from IntA form the substrate of its control mechanism and are unlikely to change substantially over the lifespan of the animal. This stable relationship between IntA activity modulation and movement kinematics could facilitate precise movement by allowing Purkinje firing patterns direct access to a controller, before, during and after learning. This arrangement would simplify the learning rules for Purkinje neurons, since the converse would require that Purkinje neurons learn not only novel sensorimotor associations but also a plastic relationship between cerebellar nuclear activation and motor output. Instead, the relatively stable relationship demonstrated here unburdens the cerebellar cortex by allowing Purkinje cell activity direct access to a directionally-consistent kinematic controller.

The properties of kinematic control exerted by IntA suggest the existence of a bidirectional, continuous neural code, where firing rates scale monotonically with the amplitude of effect on movement kinematics. When we decreased the optical power of stimulation, we found concomitantly smaller amplitude decreases in outward and upward velocity, indicating that endogenous fluctuations in cerebellar nuclear activity may alter movement kinematics with a particular direction and amplitude. Importantly, this interpretation relies on conceiving of optogenetic excitation and inhibition experiments as a broad examination of the underlying neural code, as opposed to a more limited view as binary gain-or loss-of-function manipulations(Jazayeri and Afraz, 2017). Our closed loop system enabled exploration of a wide dynamic range of neural modulation at a particular behavioral context (i.e. a specific point in reach), allowing us to demonstrate that modulation of IntA in both directions exerts meaningful effects on downstream structures. Previous work argued for a predominant role of Purkinje disinhibition in producing movement (Witter et al., 2013; Lee et al., 2015), implying that cerebellar nuclear activity has a unidirectional relationship to kinematics. Our finding that inhibition of IntA during reach increased outward and upward velocity provides a mechanism by which increases in Purkinje firing, as opposed to solely decreases, could exert kinematic control. Notably, in vivo recordings of Purkinje neuron activity during motor behavior often finds increases in rates (Pasalar et al., 2006; Herzfeld et al., 2015), thus our findings have broad relevance for continuous control of movements. Moreover, the bidirectional and monotonic nature of the functional relationship endows IntA with the ability to precisely modify ongoing limb movements, in strong agreement with the well-known deficits associated with cerebellar damage.

In order to more broadly characterize IntA’s causal relationship to movement kinematics, we expanded our experiments to other unique landmarks during reach. We found that, regardless of initial position or velocity of the limb at the time of stimulation, the direction of the kinematic effect observed was consistent, revealing another layer of stability for the translation of cerebellar output into movement. Quantitative analysis of the amplitude of kinematic effects revealed differences across stimulation locations, but crucially did not support the hypothesis that IntA controls reach progression, since identical kinematic effects were seen on outreach and return. Instead, we argue that cerebellar kinematic commands are added onto the underlying motor command structure, in general agreement with functional studies of eye movements (Skavenski and Robinson, 1973; Ramachandran and Lisberger, 2006). The small context dependence of kinematic effect amplitude could result from intrinsic nonlinearities in either the biomechanical properties of the limb or the biophysical properties of downstream neuronal circuits. One difficulty in providing a more precise mechanistic description of cerebellar contributions to the underlying motor plan is the lack of consensus about the nature of the underlying code or control mechanisms in other motor control regions (Shenoy et al., 2013; Georgopoulos and Carpenter, 2015; Omrani et al., 2017). Because IntA targets multiple downstream motor control regions important for reaching, future experiments testing how cerebellar output contributes to the motor command structure of efferent targets may help elucidate the precise mechanisms of cerebellar kinematic control demonstrated here.

### Implications of kinematic control by IntA for cerebellar theory

Previous studies attempting to determine the functional role of the cerebellar nuclei during movement have mainly focused on reflexive behaviors(Heiney et al., 2014b; Kodama and du Lac, 2016) or used chemical lesions lasting orders-of-magnitude longer than a single movement (Milak et al., 1997; Mason et al., 1998; Cooper et al., 2000; Martin et al., 2000; Low et al., 2018). By replicating the clinical finding of dysmetria, the latter category of studies importantly identified IntA as a critical locus for forelimb control, but the nature of the kinematic effects observed varied widely across experiments, and included both hypo- and hypermetric effects (Cooper et al., 2000; Low et al., 2018). In this study, our ability to track reach kinematics in real time, coupled with the temporal precision afforded by optogenetics, allowed us to control the behavioral context of neural manipulation to an unprecedented degree (i.e. a specific point in reach), and in doing so, define the acute functional importance of cerebellar output to ongoing limb control.

In addition, we demonstrate that IntA activity can both actively and precisely shape purposive reaching movements. While correlations between cerebellar neural activity and movement parameters are well documented (Burton and Onoda, 1977; Fortier et al., 1989; Coltz et al., 1999; Pasalar et al., 2006; Heiney et al., 2014a; Herzfeld et al., 2015; Chen et al., 2016), it has remained difficult to distinguish real-time contributions from movement initiation and/or planning. Our evidence that cerebellar output can sculpt kinematics in real time comports well with some viewpoints of cerebellar nuclear function during delayed eyeblink conditioning, where precisely timed cerebellar activity modulates eyeblink kinematics (Heiney et al., 2014b). Moreover, cerebellar-mediated eyeblink conditioning at different time intervals appears to modulate the kinematics of eyelid closure, as opposed to simply altering eyeblink timing, implying that the substrate of cerebellar control through associative learning may be movement kinematics itself (Chettih et al., 2011). In addition, our observation that the amplitude of kinematic effect depends significantly on behavioral context (i.e. between reaching and quiescence) agrees well with the view that cerebellar output sculpts ongoing motor commands generated elsewhere, instead of initiating them de novo (Eccles et al., 1967).

Though the existence of a continuous code for kinematic control in IntA is consistent with many different algorithmic theories of cerebellar function (Miall and Wolpert, 1996; Wolpert et al., 1998), it significantly informs the properties through which any such implementation is executed. One possibility is that cerebellar output from IntA encodes a prediction of future kinematic state (e.g. velocity). Under this paradigm, the bidirectional velocity changes observed under excitation and inhibition would be due to altered predictions that are misaligned relative to the actual state of the arm. For example, increased population activity in IntA during outreach might inform downstream structures that the limb is predicted to be moving too fast in the outward direction, resulting in subsequent motor commands that decrease outward velocity. Another possibility is that a comparison between prediction and current state happens at the level of the cerebellar nuclei themselves (Brooks et al., 2015; Herzfeld and Shadmehr, 2016). In this case, learned Purkinje predictions would be integrated with mossy fiber collaterals that signal current state, summing together to produce an error-corrective motor command. Stimulation of IntA during reach would be interpreted as an injection of an exogenous motor ‘correction’ for an error that does not actually exist. Integration of our findings with future work elucidating the encoding properties of IntA motor output neurons could provide fundamental insight into the algorithms executed by cerebellar circuitry.

## Methods

### Subjects

Adult (>8 weeks old) wild-type C57/Bl6 mice of either sex were used in all experiments. Animals were housed on a 12:12 light-dark cycle with ad libitum access to food and water except during behavioral training and experimentation (described below). All procedures were in accordance with NIH guidelines for the care and use of laboratory animals and were approved by the University of Colorado Anschutz Institutional Animal Care and Use Committee and Institutional Biosafety Committee.

### Behavior

Animals were trained on a skilled reach task described previously(Whishaw, 1996; Azim et al., 2014). Briefly, animals were food restricted to 80-90% their initial body weight and monitored daily for weight gain or loss as well as signs of distress. They were then accommodated to the behavioral arena, and subsequently trained to reach for 20 mg pellets of food (BioServ #F0163). The behavioral arena consisted of a custom plexiglass box with a 0.9 cm opening providing access to a cylindrical pedestal that held a food pellet 1 cm away, 2 cm from the bottom of the behavioral arena and slightly left-of-center to encourage reaching with the right hand. The pellet pedestal (0.5 cm diameter) was designed to avoid physical interference with outreach or return limb reach trajectories and required the animals to skillfully grab the pellet to retrieve it. On training day 1, the pellet location was moved close enough to the arena opening to be retrieved with the tongue, and was slowly moved farther away on subsequent days to encourage reaching. All animals learned to accomplish the task using the right hand (n = 7; range of 1 to 10 days of training). Sessions lasted until 30 pellets were successfully retrieved or 20 minutes were spent in the arena, whichever came first. Success was defined as bringing the pellet into the behavioral arena. Animals were considered ‘trained’ and ready for experimentation when they could successfully retrieve 30 pellets in a 20 minute behavioral session.

### Real-time kinematic tracking and closed-loop system

Paw position was monitored by an infrared-based real-time motion-capture system consisting of five 120 frames-per-second cameras (Optitrack Slim3U Camera board with Motive), each with a 10 mm focal length lens (Edmund Optics). Custom built camera mounts allowed stable camera positioning day-to-day and three-degree-of-freedom control over positioning. Cameras were mounted with infrared LED ring arrays and light level was controlled via digital power supply (Hewlett Packard). Camera position and orientation was optimized to capture the mouse’s paw movement throughout the extent of the reach, resulting in a capture volume of approximately 4 cm^3^. Four cameras were used for online kinematic tracking, and one was used for reference video. 1.5 mm diameter retroreflective markers (B&L Engineering) were used for both camera calibration and tracking of mouse reach kinematics. Camera calibration was conducted in Optitrack Motive software with a custom built calibration wand (4 and 8 mm marker spacing) and ground plane (10 and 15 mm marker spacing). The spatial origin of the reach capture volume was located approximately 1.6 cm inside the behavioral arena, measured from the front plexiglass wall, and approximately 2.0 mm left-of-center from the reach opening (in line with the pellet target location). Calibration procedures were identical throughout the study, and the system was recalibrated as necessary to maintain accurate detection. After calibration, Motive reported mean spatial triangulation errors of less than 0.05 mm throughout all experimental sessions.

For kinematic tracking, marker detection thresholds were set to minimize spurious detection of non-marker objects (e.g. the mouse’s eye or snout). Real-time tracking of paw position was conducted by Motive and streamed into Matlab (R2015b; RRID: SCR_001622) for processing, with a tracking latency of less than 1 ms. A custom-written Matlab program detected when the paw crossed a user-defined boundary and sent a ‘go’ signal to an Arduino microcontroller, pre-programed to drive a laser via TTL pulses. We modified an open-source C++ dynamic link library(Stavropoulos, 2015) to facilitate low-latency communication between Matlab and Arduino (0.5 ± 0.1 ms (mean ± standard deviation) reflection latency). Combined with the camera frame rate (120 Hz), this system supports a closed-loop latency of 8.5-10 ms. The three-dimensional reach position data was saved in Matlab along with the time of stimulation and reach success. Reach success was monitored manually by the experimenter through a keypress function in Matlab.

### Surgical procedures and histology

All surgical procedures were conducted under Ketamine/Xylazine anesthesia. The stereotaxic location of Anterior Interposed nucleus (IntA) was targeted as -1.95 mm posterior, 1.6 mm lateral, and 2.3 mm ventral from lambda. Pressure viral injections were performed with a pulled glass pipette. Approximately 150 nL of virus was injected unilaterally into the right IntA, ipsilateral to the paw used for reaching, over the course of approximately 1 minute. A minimum of four weeks was allowed for expression before optogenetic stimulation experiments. For ‘ChR2’ experiments, we used AAV2-hSyn-hChR2(H134R)-mCherry; for ‘Arch’ experiments, we used AAV2-hSyn-eArch3.0-EYFP (UNC Vector Core). Optical fibers (105 µm core diameter, ThorLabs) attached to a ceramic ferrule (1.25 mm, ThorLabs), polished to efficiency >90%, were implanted so that the tip of the optical fiber rested 0.1 mm above the injection site. The ceramic ferrule was affixed to the skull using luting (3M) and dental acrylic (Teet’s cold cure). After behavioral experiments were completed, animals were sacrificed according to standard procedures via pentobarbital overdose, transcardially perfused with 4% paraformaldehyde, and processed for histological analysis as described previously(Beitzel et al., 2017). Injection sites and fiber implant tracts were visually inspected on an upright epifluorescent microscope (Zeiss) by two independent observers.

### Optogenetics

Stimulation for ChR2 experiments was obtained via activation of a 473 nm diode laser (SLOC) with 2 ms pulses at 100 Hz, 50 ms pulse train duration. During kinematic closed-loop experiments, a custom-built patch cord (approx. 1.0 m) connected the laser to the implanted ferrule and was secured with a ceramic sleeve connector (ThorLabs). For ChR2 experiments, power was set to between 0.5-1.0 mW, measured at the end of the patch cord. For experiments shown in Fig. 3 that use light power as an experimental variable, powers between 0.05-2.0 mW were used, with ranges varying individually between animals. Stimulation for Arch experiments was obtained via activation of a 561 nm diode laser (SLOC) with a single 50 ms pulse with light power at 5.0 mW. Light power levels were calibrated daily. The irradiance at the tip of the 105 µm diameter optical fiber ranged from approximately 5.0 mW/mm^2^ (0.05 mW) to 230 mW/mm^2^ (2.0 mW) for ChR2 experiments, and was 580 mW/mm^2^ (5.0 mW) for Arch experiments.

Stimulation occurred on a pseudorandom 25 percent of reaches to avoid anticipation. Stimulation occurred in closed-loop based on a kinematic landmark defined prior to the experimental session. All experiments used a positional threshold in the outward direction near the point of maximum outward reach velocity as the kinematic landmark (Supplementary Fig. 1), unless otherwise noted in experiments directly manipulating stimulation location. This main kinematic landmark (‘Middle’ in Fig. 4) was a vertical plane 1.6 cm from the ‘origin point’, defined during calibration, which corresponds to the opening in the front acrylic wall of the behavioral arena. The ‘Early’ kinematic landmark was located 1.4 cm from the origin, while the ‘Late’ kinematic landmark was located 2.1 cm from the origin. Stimulation during return was performed at the position of the ‘Late’ kinematic landmark following completion of outreach, as measured by a direction reversal. Stimulation during quiescence and/or feeding used identical stimulation parameters, except that stimulation was triggered every 10 seconds as the animals rested or ate food pellets placed inside the behavioral arena.

### Kinematic analysis

Data were analyzed with custom-written functions in Matlab. Time-stamped three-dimensional paw position data (reported as values relative to the spatial origin set during calibration) were processed in several stages to segment data into individual reaches. First, we corrected for spurious object detection (<3% of data frames detected more than one marker) by conducting a nearest-neighbor analysis, linking marker positions across frames to differentiate the marker from other objects. Next, we filtered continuous kinematic data captured throughout the behavioral session, including while the animal freely moved in the behavioral arena and performed reaches. Between reaches, the marker could become hidden from view, thus to avoid artefacts from erroneously linking distant marker data, we linearly interpolated missing points prior to filtering. After applying a 2^nd^ order 10 Hz cutoff Butterworth fitler(Yu et al., 1999), we removed the interpolated frames, which resulted in continuous filtered marker position of data captured during the experiment.

To segment reaches from the continuously collected kinematic data, we extracted marker positions beyond a positional threshold unique to reaching, located just beyond the reach opening of the arena. To ensure that we captured the entirety of the reach, we included marker data that continuously moved in the outward direction prior to the paw reaching the positional threshold. The inverse criterion was applied to the end of the reach to obtain full reach trajectories.

Reach velocity was calculated as the numerical gradient of reach position data in each dimension. Speed measurements were calculated as the three-dimensional Euclidean distance in the velocity data. We defined ‘outreach’ as the period from reach initiation (defined above) up to the first point at which the paw moved back towards the body. To analyze positional effects, we clipped reach data at the first local maximum point in the outward direction. Outreach duration was calculated for this clipped data set for unstimulated reaches. To produce average reach velocity profiles for unstimulated or stimulated reaches, we first time-interpolated the velocity data at 10 ms intervals over a 400 ms window aligned to time of stimulation. For unstimulated reaches, the alignment point was chosen as the point at which stimulation would have occurred during that behavioral session. Average velocity at each time point is reported as the mean of the population of reaches under consideration. Since the reach data traces could have different lengths, standard error was calculated on a point-by-point basis for each average. Two-dimensional plots of velocity effects were generated by subtracting the average unstimulated and stimulated velocity values in each dimension. We calculated the latency to maximum effect size on a per animal basis by finding the peak Euclidean difference in average velocity between stimulated and unstimulated reaches. The amplitude of the effect was reported for each animal at its calculated latency of maximum effect.

To analyze the distribution of velocity values within the population of unstimulated or stimulated reaches, we generated a two-dimensional histogram over the range of possible velocity values over the 400 ms time window. These histograms were normalized to the total number of data points per condition to obtain a relative ‘prevalence’ of velocity values in each bin. These values were then subtracted on a bin-by-bin basis (Stimulated – Unstimulated) to visualize differences in the likelihood of velocity measurements between unstimulated and stimulated reaches as populations (Fig. 3B), displayed as heatmaps.

Nonparametric statistical tests were used for all analyses. All kinematic data are displayed as mean ± standard error unless otherwise noted. Reach population sizes for both unstimulated and stimulated reaches are organized on a per animal basis in Supplementary Table 1. To test for statistical difference in velocity between Unstimulated and Stimulated reaches, the point of maximum difference between the two average velocity profiles was used. For continuous z-score analysis, a Wilcoxon rank-sum test was conducted on the time interpolated averages and the associated z-score was calculated at each time point. The first time point to surpass a z-score of 2.5 was defined as the latency to divergence of the two velocity averages. For visualization, the color bar displaying the z-score was linearly interpolated.

### Recordings

Mice injected with AAV2-hSyn-hChR2(H134R)-mCherry to the cerebellar nuclei were implanted with an optetrode drive(Anikeeva et al., 2011). Four 0.012 mm NiCr wire tetrodes affixed to a 105 µm core optical fiber were attached to an electrode interface board (Neuralynx) and 3D-printed movable drive. Tetrodes were electrochemically plated with gold solution to an impedance of 250 kΩ. Tetrodes were positioned 0.5mm below the tip of the optical fiber. The optical fiber/tetrode bundle was implanted 0.3 mm above IntA (-1.95 mm posterior, 1.6 mm lateral, and 2.0 mm ventral from lambda) and affixed to the skull using luting (3M) and dental acrylic (Teet’s cold cure). Neurons were recorded as the animal rested or ate food pellets placed inside the behavioral arena, and optogenetic stimulation was applied as described for behavioral experiments (1.0 mW, 100 Hz, 2 ms pulses, 50 ms train). The drive was incrementally lowered by 0.05 mm the day before each recording session. Recordings were conducted with a Cereplex M digital headstage connected to a Cereplex Direct acquisition system (Blackrock Microsystems), which also monitored optical stimulation timing via TTL. Blackrock Offline Spike Sorter software was utilized to sort individual units from noise, with spike amplitudes being at least a factor 4.0 larger than noise root mean square values. Spike times were subsequently imported into Matlab for analysis. Two-sided paired t-tests were conducted to compare firing rates before (100 ms) and during (50 ms) optogenetic stimulation for each cell (n = 6).

Mice injected with AAV2-hSyn-eArch3.0-EYFP to the cerebellar nuclei (>4 months old) were deeply anaesthetized with isofluorane and transcardially perfused with warm (36?°C) ACSF containing (in mM): 123 NaCl, 3.5 KCl, 26 NaHCO_3_, 1.25 NaH_2_PO_4_, 1.5 CaCl_2_, 1 MgCl_2_, 10 glucose and equilibrated with 95/5% O_2_/CO_2_. Mice were then rapidly decapitated and the brains removed into ACSF (36?°C). Slices (300?μm thick) were cut on a Vibratome (Leica VT 100S) and incubated in warmed (37?°C), oxygenated ACSF for at least 1?h before recording. Cerebellar slices were transferred to a recording chamber perfused continuously with warmed (30–32?°C), oxygenated ACSF at a flow rate of 2–4?ml/min. Slices were visualized with infrared differential interference contrast microscopy and fluorescence (Zeiss AxioExaminer) and recordings were made from neurons expressing GFP. Borosilicate patch pipettes were pulled to tip resistances of 4?MΩ and filled with an internal solution containing (mM): 130 K-gluconate, 2 Na-gluconate, 6 NaCl, 10 HEPES, 2 MgCl2, 0.1 or 1 EGTA, 14 Tris-creatine phosphate, 4 MgATP, 0.3 Tris-GTP and 10 sucrose. A 105 um core optical fiber was positioned over the slice and coupled to a 561 nm diode laser (SLOC). Recordings were made in the on-cell configuration to monitor spontaneous firing. 50 ms light pulses were delivered 1/s after achieving a recording collecting over 100 sweeps/neuron (n = 5). Two-sided paired t-tests were conducted to compare firing rates before (100 ms) and during (50 ms) optogenetic stimulation for each cell.

## Data availability

Data analyzed in this study is available upon written request to corresponding author.

## Code availability

Code is available upon written request to corresponding author.

## Acknowledgements

We thank Drs. Gidon Felsen, Matthew Kennedy, Joel Zylberberg, Cristin Welle, and members of the Person lab for their insightful comments on the manuscript. Samantha Lewis provided technical assistance. Light microscopy was performed in the University of Colorado Anschutz Medical Campus Advance Light Microscopy Core. Optogenetics support was provided by the University of Colorado Optogenetics and Neural Engineering Core. Both cores are supported in part by Rocky Mountain Neurological Disorders Core Grant Number P30NS048154 and by NIH/NCRR Colorado CTSI Grant Number UL1 RR025780. This work was supported by National Research Service Award Individual Predoctoral fellowship (F31) NS103328 to M.I.B; the McKnight Foundation, Klingenstein Foundation; NSF 1749568 to A.L.P.

## Contributions

M.I.B. and A.L.P. designed and conceived of the experiments. M.I.B. built the kinematic closed-loop system, conducted experiments, and performed analyses. A.L.P. oversaw all aspects of experiments. M.I.B. and A.L.P. wrote the paper.

## Conflict of Interest

None.

## Competing Interests Statement

The authors declare no competing interests.

